# *Tie that binds* - Hill-Robertson Interference can produce signals of sexual antagonism even in the absence of sexually antagonistic selection

**DOI:** 10.1101/2025.01.08.632022

**Authors:** Manas Geeta Arun

**Affiliations:** The University of Edinburgh

## Abstract

Sexually antagonistic (SA) selection on a trait with a shared genetic basis between the sexes is thought to lead to Intralocus Sexual Conflict (IaSC). A common way of measuring IaSC involves characterising the joint distribution of breeding values of male and female fitness using the intersexual additive genetic correlation for fitness (*r*_*mf*,*W*_). However, there is considerable variation in the empirical estimates of *r*_*mf*,*W*_ - with estimates ranging from significantly negative (suggestive of strong IaSC) to significantly positive (suggesting an absence of IaSC) - even among closely related populations of the same species. Here, we investigate whether SA selection is even necessary for *r*_*mf*,*W*_ to be negative. We first highlight that the sign of *r*_*mf*,*W*_ depends on a term that describes direct sex-specific selection, and another that describes indirect selection mediated by linkage disequilibria (LD). We show that *r*_*mf*,*W*_ can be negative if (1) selection is perfectly sex-limited - such that every locus affects the fitness of one sex only - and (2) the LD between closely-linked male-beneficial and female-beneficial alleles is on average negative as predicted under Hill-Robertson Interference (HRI). We test this idea using individual-based simulations. Our results suggest that populations evolving under purely sex-limited selection exhibit a prominent negative bias in *r*_*mf*,*W*_ that gets strengthened by a reduction in the recombination rate. Therefore, HRI can drive *r*_*mf*,*W*_ to be negative even in the absence of SA selection suggesting that negative estimates of *r*_*mf*,*W*_ should not automatically be considered as evidence of IaSC.

## 2 Introduction

Males and females of many species have startlingly different morphology, behaviour, and physiology (Kappeler *et al*., 2023). At the same time, they share - with important exceptions such as sex chromosomes and mitochondria - almost the entirety of the population’s gene pool with each other. In some sense, the sexes can be thought of as sub-populations that are exposed to two very different selective environments with unrestricted gene flow between them (Connallon & Clark, 2010). Therefore, selection experienced by one sex can interfere with adaptation in the other sex. This selective interference between the sexes can result from direct or indirect selection. First, loci that affect fitness in one sex may have (potentially different) fitness consequences when expressed in the opposite sex. If a locus has opposite fitness effects in the two sexes, the resulting sexually antagonistic (SA) selection has the potential to result in a polymorphic equilibrium (Owen, 1953; Parsons, 1961; Haldane, 1962; Kidwell *et al*., 1977) - albeit only when the strength of selection is unusually high. Alternatively, selective interference between the sexes can be subtle, driven by indirect selection. If loci that affect fitness in one sex are in linkage disequilibrium (LD) with a second set of loci that affect fitness in the other sex, selection experienced in one sex can lead to indirect selection in the other sex. However, both from a theoretical and empirical point of view, the role of indirect selection in mediating selective interference between the sexes remains under-explored.

Selective interference between the sexes has primarily been investigated using the quantitative genetic framework of Intralocus Sexual Conflict (IaSC) (Bonduriansky & Chenoweth, 2009; Van Doorn, 2009). In this framework, if a trait common to both sexes - specifically, a trait with a positive intersexual additive genetic correlation (*r*_*mf*_) - is subjected to SA selection, the population as a whole may remain stuck in a maladaptive equilibrium where neither sex is at its fitness optimum. IaSC is thought to be resolved by the evolution of sex-limited genetic architecture such as sex-specific gene expression (Con-nallon & Clark, 2010), sex-specific genetic imprinting (Day & Bonduriansky, 2004; Spencer & Clark, 2014), sex-specific dominance patterns (Spencer & Priest, 2016), etc. While the mathematical foundation of IaSC was laid more than four decades ago (Lande, 1980), empirical investigations began only after the turn of the century. A number of studies have detected signatures of IaSC on traits such as locomotory activity (Long & Rice, 2007) and immunocompetence (Vincent & Sharp, 2014) in *Drosophila melanogaster*, leaf area in *Silene latifolia* (Delph *et al*., 2011), bill colour in *Taeniopygia guttata* (Price & Burley, 1993), multivariate life-history syndrome in *Callosobruchus maculatus* (Berger *et al*., 2016), among others. Nevertheless, organisms have an enormously large number of fitness-determining traits that interact with each other in complicated ways. Identifying candidate traits involved in IaSC, investigating sex-specific genetic architecture and sex-specific selection on those traits can be a challenging undertaking. To get around this problem, a number of studies have attempted to detect averaged signals of IaSC at the level of overall fitness. This typically involves characterising the joint distribution of male and female breeding values for fitness using metrics such as the intersexual additive genetic correlation for fitness (*r*_*mf*,*W*_) (Bondurian-sky & Chenoweth, 2009). If IaSC is intense, such that there is predominantly SA selection on shared genetic variation between males and females, *r*_*mf*,*W*_ would be negative. On the other hand, if sexually concordant (SC) selection on shared genetic variation between the sexes predominates, *r*_*mf*,*W*_ would be positive. Lastly, if the sexes have fully sex-limited genetic architectures, such that loci affect the fitness of one sex only, *r*_*mf*,*W*_ would be 0 (see Figure 1 in Bonduriansky & Chenoweth (2009)).

**Figure 1:**
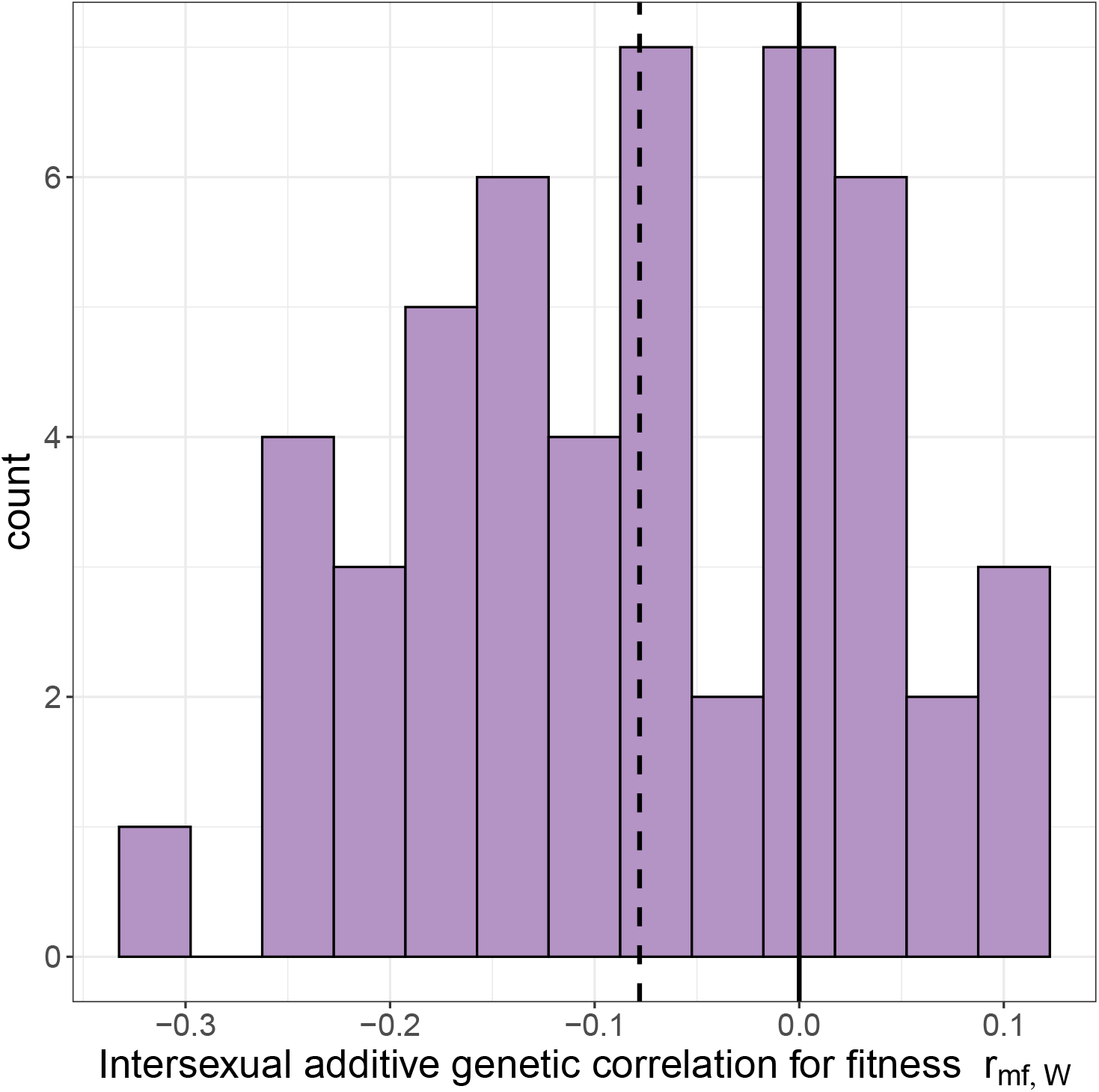
The distribution of *r*_*mf*,*W*_ after the population was allowed to evolve for 25000 generations. The dashed vertical line is the mean of the distribution and the solid line indicates 0. In these simulations, the map length of the simulated region was 0.1 M and the mutation rate was 7 × 10^*−*7^.

The first decade of this century was characterised by a string of studies reporting negative estimates of *r*_*mf*,*W*_ in organisms such as fruit flies *D. melaogaster* (Chippindale *et al*., 2001; Innocenti & Morrow, 2010) and *Drosophila serrata* (Delcourt *et al*., 2009), the field cricket *Allonemobius socius* (Fedorka & Mousseau, 2004), red deer *Cervus elaphus* (Foerster *et al*., 2007), etc. However, there is considerable variation in some of the more recent estimates of *r*_*mf*,*W*_, with many studies reporting estimates that were either not distinguishable from 0 (Berger *et al*., 2014; Collet *et al*., 2016; Martinossi-Allibert *et al*., 2018; Wolak *et al*., 2018; Ruzicka *et al*., 2019; Collet & Sztepanacz, 2022) or estimates that were, in fact, significantly greater than 0 (Berger *et al*., 2014; Geeta Arun *et al*., 2022; Singh *et al*., 2023; Wong & Holman, 2023). More strikingly, even estimates from closely related populations of the same species, such as the various replicates of the laboratory adapted *LH*_*m*_ population of *D. melanogaster* - all maintained in nearly identical laboratory environments - have varied from being significantly negative (Chippindale *et al*., 2001; Innocenti & Morrow, 2010; Collet *et al*., 2016) to indistinguishable from 0 (Collet *et al*., 2016; Ruzicka *et al*., 2019) to significantly positive (Geeta Arun *et al*., 2022). One explanation for these patterns is that sex-specific genetic architecture is fairly malleable such that the strength of IaSC evolves rapidly between even closely related populations of the same species. This, however, rests on the assumption that *r*_*mf*,*W*_ is a reliable indicator of the strength of IaSC.

In a landmark study, Connallon & Matthews (2019) developed expressions for *r*_*mf*,*W*_ as a function of (1) the strength of directional selection experienced by the two sexes, (2) the strength of intersexual genetic correlations for traits under selection, and (3) the degree of SA selection measured as an angle between selection vectors in males and females. Their results suggest that SA selection is not sufficient for *r*_*mf*,*W*_ to be negative, which also requires that “the magnitude of the displacement of each sex from its optimum … must be large relative to the magnitude of the shared genetic variance in the set of traits” (Connallon & Matthews, 2019). As a corollary, this also means that a positive *r*_*mf*,*W*_ does not necessarily preclude IaSC. Nevertheless, their model did seem to suggest that SA selection (such that the angle between the selection vectors in males and females is between 90° and 180°) is indeed a necessary condition for *r*_*mf*,*W*_ to be negative. An important caveat in the multivariate models by Connallon & Matthews (2019) was the assumption that different traits in the model were not genetically correlated with each other and that correlational selection was absent. In other words they assumed that there is no linkage disequilibrium (LD) between loci that code for different traits. However, in populations of finite size, selection at one locus can interfere with selection at a closely linked locus resulting in the build-up of negative LD between closely linked beneficial mutations (Hill & Robertson, 1966; Felsenstein, 1974; Roze & Barton, 2006). Crucially, Hill-Robertson Interference (HRI) is agnostic to the sex in which selection operates and only predicts the build-up of negative LD between closely linked mutations that are beneficial when averaged over both sexes.

In this study we relax the assumption of linkage equilibrium and reexamine whether IaSC and SA selection are even necessary for *r*_*mf*,*W*_ to be negative - especially in the context of indirect selective effects between males and female. Specifically, we focus on purely sex-limited selection, such that each locus affects the fitness of one sex only. In the framework of IaSC this implies fully resolved conflict. In the context of the model by Connallon & Matthews (2019) this would mean that the selection vectors in males and females are perfectly orthogonal. If a locus that affects female fitness only is closely linked to another locus that affects male fitness only, HRI predicts the build-up of negative LD between beneficial alleles at the two loci. First, using a brief mathematical argument we show that this can lead to negative *r*_*mf*,*W*_ even in the absence of SA selection. Next, we test this idea by performing multilocus individual-based simulations implemented using SLiM 4.2 (Haller & Messer, 2023).

## 3 A closer look at *r*_*mf*,*W*_

In this section we investigate how indirect selection between the sexes can drive *r*_*mf*,*W*_ to be negative when selection is purely sex-limited. Consider a population of diploid individuals where male and female fitness variation is determined by *N*_*L*_ biallelic segregating loci. Let *p*_*i*_ be the frequency of the reference allele at locus i. An individual’s genotype at each locus can be described in terms of the proportion of copies of the reference allele at that locus, either 1, 0.5, or 0, depending on whether the individual is a homozygote for the reference allele, or a heterozygote, or a homozygote for the non-reference allele at that locus respectively. Let **L** be the genetic variance-covariance matrix for the population obtained y computing the covariance of genotypic states at various loci over individuals. Under random mating, the diagonal elements of **L** represent genetic diversities, *p*_*i*_(1*− p*_*i*_)/2, at each locus, while the off-diagonal elements represent signed linkage disequilibria (LD) between pairs of loci. Let *D*_*i*,*j*_ be the LD between locus i and locus j; note that *D*_*ij*_ = *D*_*ji*_. Fisher’s average effects for relative fitness in females and males can be represented as vectors ***α***_***f***_ and ***α***_***m***_, respectively. The intersexual additive genetic covariance for relative fitness can then be expressed as the following matrix product (similar to Equation (17) in Buffalo & Coop (2019)):

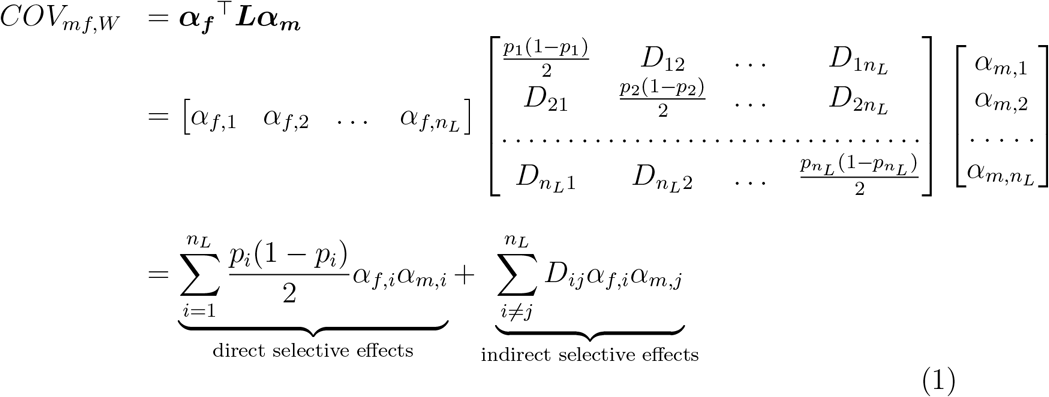

Since 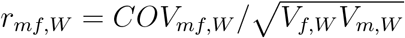 (where *V*_*f*,*W*_ and *V*_*m*,*W*_ are additive genetic variances for relative fitness in females and males respectively), *r*_*mf*,*W*_ and *COV*_*mf*,*W*_ have the same sign. Therefore, to investigate the conditions under which *r*_*mf*,*W*_ is negative we need only examine the two terms in Equation 1. If the population is at linkage equilibrium such that *D*_*i*,*j*_ = 0 for all *i≠ j*, the second term of Equation 1 is 0, i.e., indirect selective effects do not contribute to *COV*_*mf*,*W*_ . The only way *COV*_*mf*,*W*_ (and *r*_*mf*,*W*_) can be negative is if at least for some loci *α*_*f*,*i*_ and *α*_*m*,*i*_ have opposite signs. In other words, if the population is at linkage equilibrium, SA selection is indeed necessary for *r*_*mf*,*W*_ to be negative. However, in general, as a consequence of HRI the LD between closely linked loci are unlikely to be 0.

Consider a situation where selection is entirely sex-limited such that every locus affects the fitness of one sex only. This means that for no locus i, are *α*_*i*,*f*_ and *α*_*i*,*m*_ simultaneously non-zero. For example, if locus *x* is female-limited *α*_*x*,*f*_ would be non-zero, but *α*_*x*,*m*_ would be 0. The first term of Equation 1 now becomes 0, and only indirect selection contributes to *COV*_*mf*,*W*_ . Purely for the purposes of clarity, if we always chose the beneficial allele to be the reference allele, all *α*_*f*_ ‘s and *α*_*m*_’s would be positive. At the same time, with this choice of the reference allele, HRI would result in the *D*_*i*,*j*_ terms being negative for closely linked beneficial sites - irrespective of which sex these alleles are expressed in. If the LD between sufficiently many pairs of beneficial alleles were to be negative, *COV*_*mf*,*W*_ could be negative - even in the absence of SA selection.

## 4 Simulations

To investigate if purely sex-limited selection in conjunction with HRI can result in *r*_*mf*,*W*_ being negative, we performed individual-based multilocus simulations using SLiM 4.2 (Haller & Messer, 2023). We simulated a single autosome that was 1 million base-pairs long. The population size was fixed at 2500 diploid individuals throughout the simulation. Each simulation began with blank genomes. As the population evolved forward in time in SLiM’s default “Wright-Fisher” mode, two types of mutations were allowed to occur at specified rates (see below). Mutations of “type 1” affected female fitness only, while the “type 2” mutations affected male fitness only. Both type 1 and type 2 mutations were additive (dominance coefficient = 0.5) and deleterious with their selection coefficients drawn from a reflected gamma distribution with shape = 0.3 and scale = 0.05. We used an additive model for relative fitness. The breeding value for the relative fitness of an individual was calculated by summing the selection coefficients of all mutations of the appropriate type (type 1 for females and type 2 for males) present on both genomes of that individual. We added noise terms drawn from a unit normal distribution to these breeding values to obtain relative fitnesses of all individuals in the population. Following Buffalo & Coop (2020), in order to calculate the absolute fitness of an individual, we exponentiated its relative fitness. We let the population evolve for 25000 generations, after which we drew a sample of 500 individuals (250 females and 250 males) from the population and calculated the following: 1. *V*_*f*,*W*_ = ***α***_***f***_ ^*⊺*^***Lα***_***f***_, 2. *V*_*m*,*W*_ = ***α***_***m***_^*⊺*^***Lα***_***m***_, and *COV*_*mf*,*W*_ = ***α***_***f***_ ^*⊺*^***Lα***_***m***_. We performed our simulations at different recombination rates and mutation rates. First, we fixed the mutation rate at 7 × 10^*−*7^ and performed 50 replicate simulation each with the total map length of the genome being either 0.001 M, 0.01 M, 0.1 M, 1 M, or 10 M. Next, we fixed the map length at 0.1 M and performed 50 simulations each at five different mutation rates: 5 × 10^*−*7^, 7 × 10^*−*7^, 9 × 10^*−*7^, 1.1 × 10^*−*6^, and 1.3 × 10^*−*6^.

## 5 Results

Our results suggest that there was a considerable negative bias in *r*_*mf*,*W*_ at moderate levels of recombination (0.1 M) and mutation rate (7 × 10^*−*7^ per generation per site) (Figure 1) with the mean *r*_*mf*,*W*_ being -0.0780. This negative bias became stronger at lower recombination rates, but became weaker as recombination rates increased, disappearing entirely when the map length of the simulated region was 10 M (Figure 2). Furthermore, increasing the mutation rate, while keeping the recombination rate constant also resulted in an increase in the negative bias in *r*_*mf*,*W*_ (Figure 3).

**Figure 2:**
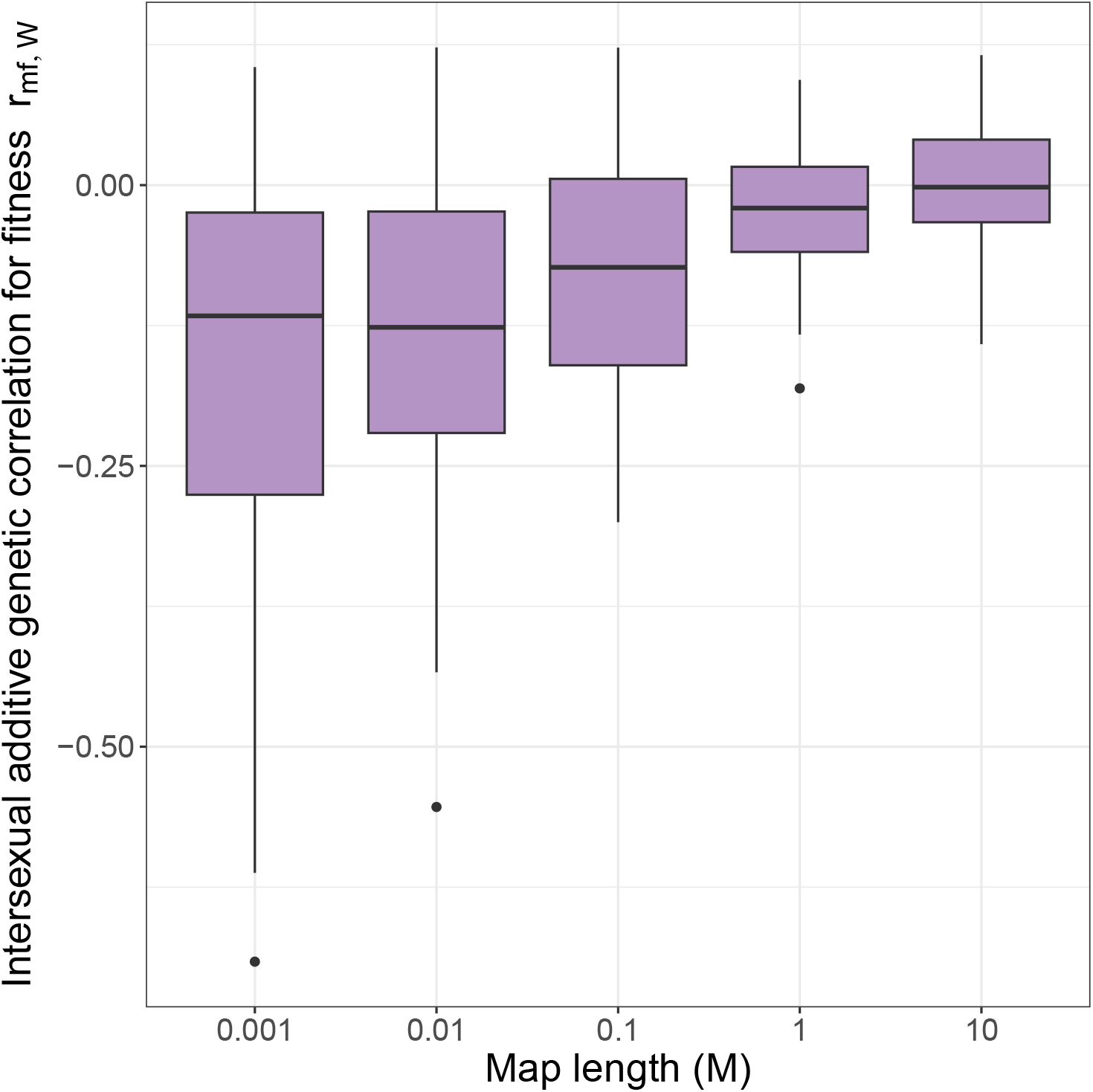
Boxplots representing the distribution of *r*_*mf*,*W*_ as a function of the map length of the region simulated. Each boxplot is based on 50 replicate simulations. In these simulations, the mutation rate was fixed at 7 × 10^*−*7^.

**Figure 3:**
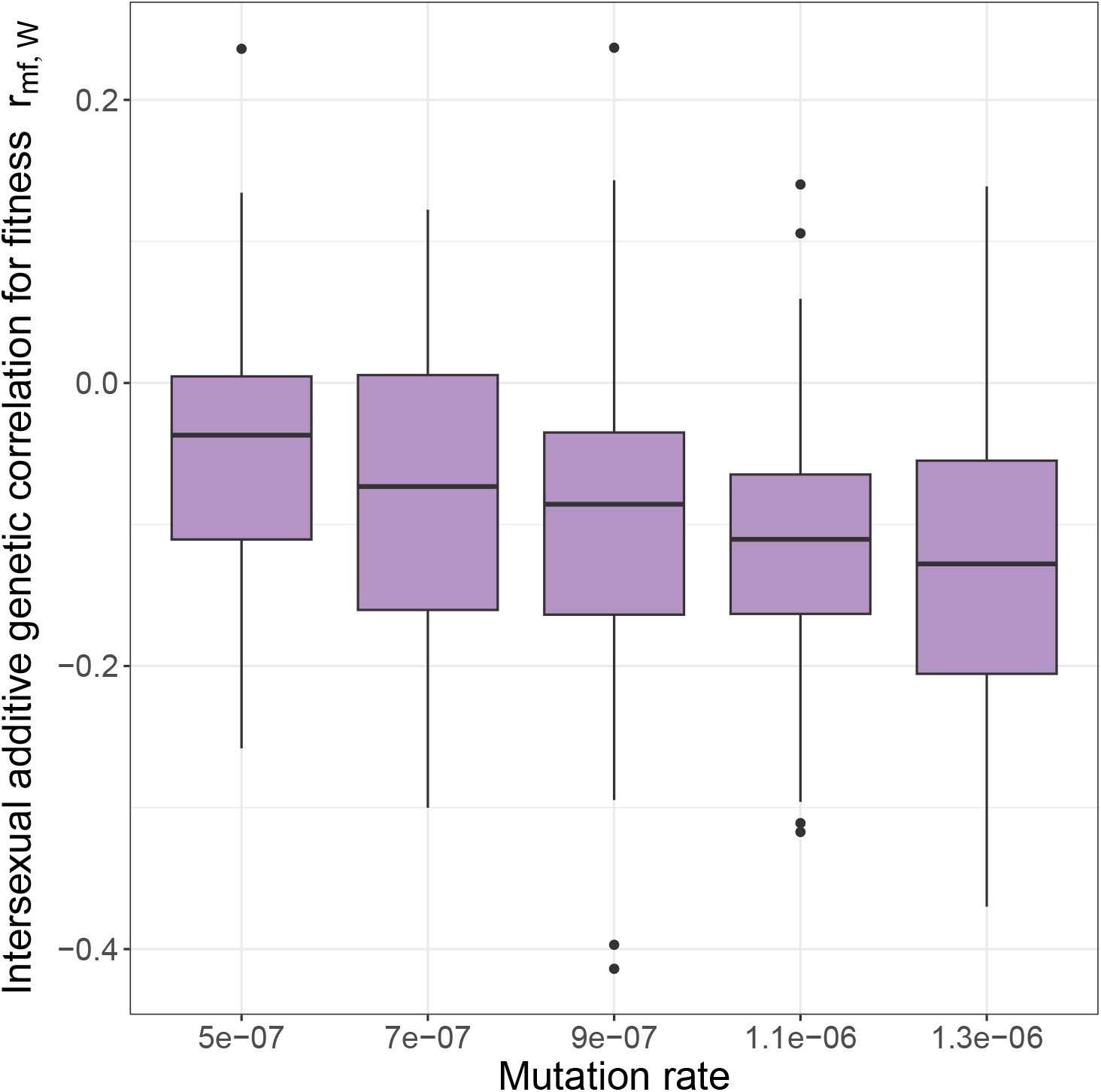
Boxplots representing the distribution of *r*_*mf*,*W*_ as a function of mutation rate. Each boxplot is based on 50 replicate simulations. In these simulations the map length of the simulated region was fixed at 0.1 M.

When the mutation rate was fixed at 7 × 10^*−*7^ both male and female additive genetic variances for relative fitness (*V*_*m*,*W*_ and *V*_*f*,*W*_) were around 0.02 in our simulations, increasing slightly with an increase in the map length of the simulated region (Figure S1A-B). The change in the total number of segregating sites with recombination rate was considerably more drastic - with the number of segregating sites going up from slightly less than 8000 to more than 13000 at 10 M (Figure S1D). When the map length of the simulated region was fixed at 0.1 M, *V*_*m*,*W*_ and *V*_*f*,*W*_ increased linearly from about 0.01 to 0.04 with mutation rate, while the number of segregating sites increased from about 6500 to over 13000 (Figure S2).

The negative bias in *r*_*mf*,*W*_ suggests that selection in one sex can negatively interfere with adaptation in the other sex. To investigate the magnitude of this interference we used the sex-specific version of Fisher’s Fundamental Theorem of Natural Selection (FTNS) (see Equation 2 in the Discussion) which expresses the rate of adaptation in one sex as the arithmetic mean of the additive genetic variance in relative fitness of that sex and the intersexual additive genetic covariance for relative fitness (*COV*_*mf*,*W*_). Since our simulations are perfectly symmetric with respect to males and females, we only present the effect of selective interference between the sexes on the rate of adaptation in females. Specifically, we measured the percentage change in the rate of adaptation in females due to selective interference between the sexes (i.e., when *COV*_*mf*,*W*_ ≠ 0) relative to an idealised situation when the selective interference between the sexes is entirely absent (i.e., when *COV*_*mf*,*W*_ assumed to be 0). This quantity can be expressed as 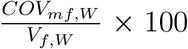 (see Discussion). Our simulations suggest that when the mutation rate was fixed at 7 × 10^*−*7^, at low map lengths, the median percentage retardation in the rate of adaptation in females was around 10%, gradually decreasing to 0 with increasing map length (Figure 4). When the map length of the simulated region was fixed at 0.1 M, selective interference led to a 5% (median) reduction in the rate of adaptation in females at low mutation rates, which increased to slightly less than 15% at higher mutation rates (Figure 5).

**Figure 4:**
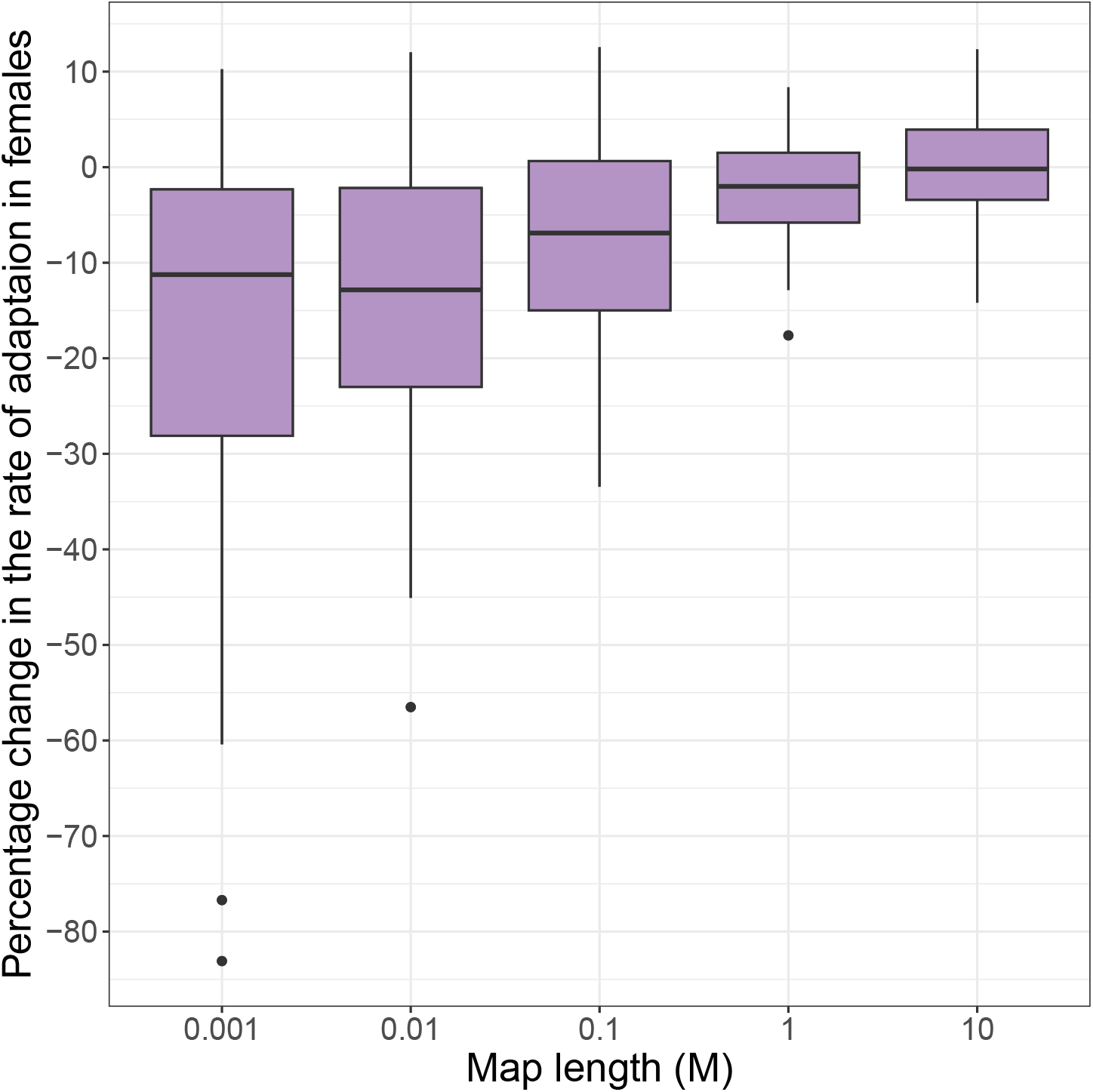
Boxplots representing the percentage change in the rate of adaptation in females due to selective interference between the sexes as a function of the map length of the simulated region. Percentage change in the rate of adaptation was calculated as 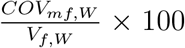. Each boxplot is based on 50 replicate simulations. In these simulations, the mutation rate was fixed at 7 × 10^*−*7^ per sit per genome.

**Figure 5:**
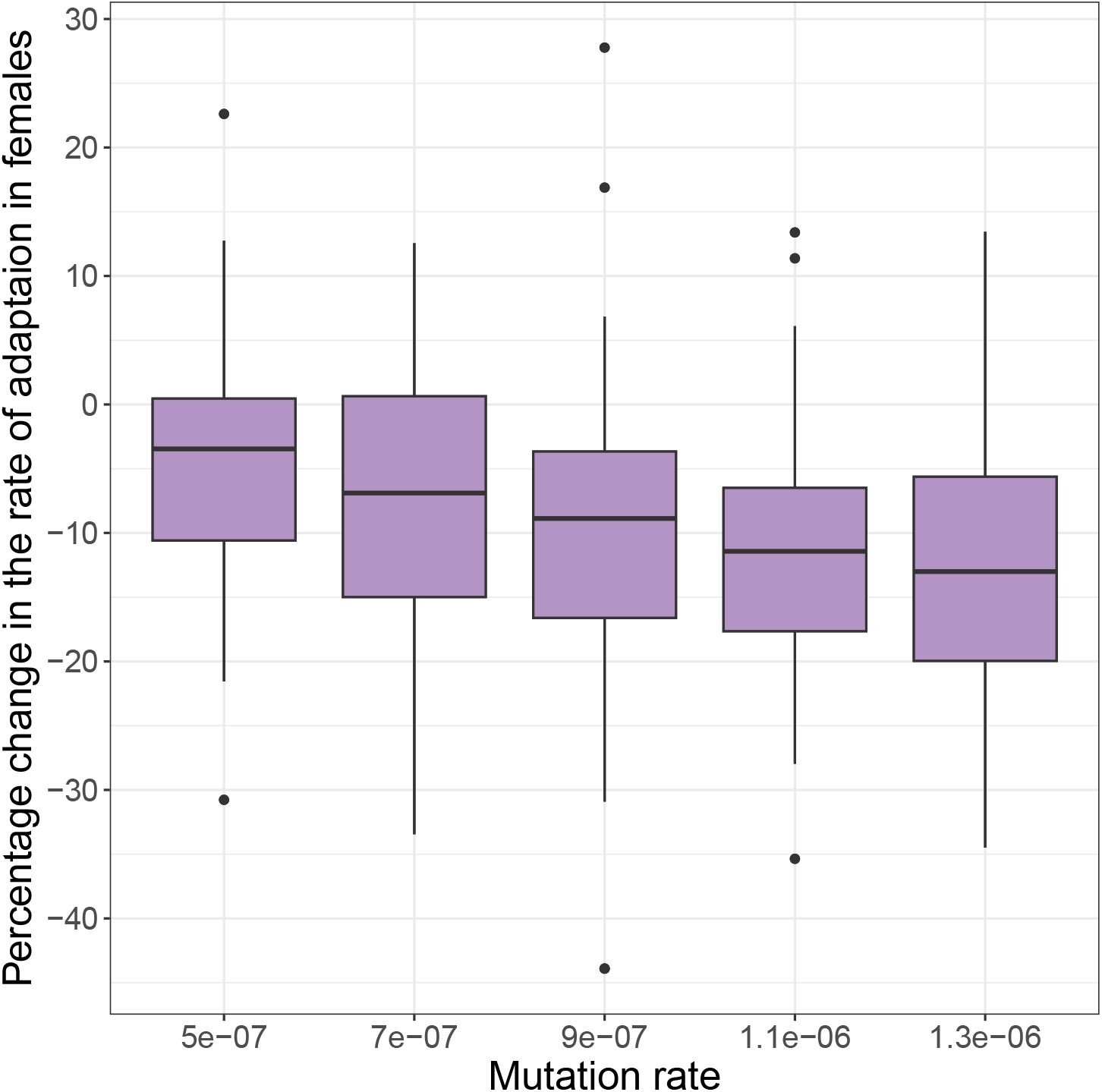
Boxplots representing the percentage change in the rate of adaptation in females due to selective interference between the sexes as a function of the mutation rate. Percentage change in the rate of adaptation was calculated as 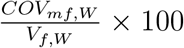. Each boxplot is based on 50 replicate simulations. In these simulations, the map length of the simulated region was fixed at 0.1 M.

## 6 Discussion

In this study we used individual-based multilocus simulations implemented in SLiM (Haller & Messer, 2023) to investigate whether purely sex-limited selection in conjunction with Hill-Robertson Interference (HRI) can result in signals of sexual antagonism. Our simulations suggest that if recombination rates are sufficiently low, purely sex-limited selection - such that each locus affects the fitness of one sex only - can result in the intersexual additive genetic correlation for relative fitness (*r*_*mf*,*W*_) being considerably negative. That this negative bias in *r*_*mf*,*W*_ was a consequence of HRI was confirmed by the observation that the *r*_*mf*,*W*_ became less negative with increasing recombination rates, disappearing entirely at high recombination rates. Consistent with this, increasing the mutation rate while keeping the recombination rate constant, such that the density of segregating sites in the genome increased, also resulted in a strengthening of the negative bias in *r*_*mf*,*W*_ . Below we discuss some major implications of these results.

### 6.1 Sexually antagonistic selection is not necessary for negative *r*_*mf*,*W*_

Empirical estimates of *r*_*mf*,*W*_ gathered over the last quarter of a century have have exhibited large variation creating uncertainty around our understanding of the ubiquity of sexually antagonistic (SA) genetic variation for fitness. Genome-wide genetic variation for fitness is likely to be a mixture of a set of loci under sexually concordant (SC) selection and another set under SA selection (Berger *et al*., 2014; Ruzicka *et al*., 2019; Singh *et al*., 2023).

Examining the properties of the joint distribution of male and female breeding values for fitness can only provide a composite picture of these SA and SC effects. Connallon & Matthews (2019) formalised this argument using a quantitative genetic model, and showed that SA selection is not sufficient for *r*_*mf*,*W*_ to be negative. It has, therefore, been argued that estimates of *r*_*mf*,*W*_ that are either positive or indistinguishable from 0 do not necessarily rule out the presence of SA genetic variation for fitness. Although not always stated explicitly in these arguments, it is generally believed that SA selection is a necessary precondition for *r*_*mf*,*W*_ to be negative. For example, a recent review suggested that “… a negative intersexual genetic correlation for fitness can indicate strong conflict over shared traits, but there may be conflict over traits linked to fitness even when the intersexual genetic correlation for fitness itself is weak or positive” (Pennell *et al*., 2024). Here, we show that this is strictly true only if the population is at linkage equilibrium. We first point out that *COV*_*mf*,*W*_ can be partitioned into a term that describes direct selection acting on loci in males and females and another term that describes indirect selective effects mediated by LD between loci that affect female and male fitness (Equation 1). If the population is at linkage equilibrium, such that indirect selective effects are absent entirely, we find that SA selection is indeed necessary for *r*_*mf*,*W*_ to be negative. In general, however, HRI predicts the build-up of substantial negative LD between closely-linked beneficial mutations. The results from our simulations suggest that in situations where the contribution of direct selective effects to *COV*_*mf*,*W*_ is small - such as when selection is purely sex-limited - negative LD between male-beneficial and female-beneficial alleles can drive *COV*_*mf*,*W*_ below 0. While our simulations are not meant to be biologically realistic, they highlight some important features of sex-specific selection. Specifically, once the assumption of linkage equilibrium is relaxed, we find SA selection is not necessary for *r*_*mf*,*W*_ to be negative. Our findings suggest that - just as a positive *r*_*mf*,*W*_ does not necessarily rule out SA fitness variation - negative *r*_*mf*,*W*_ should not automatically be considered evidence of widespread SA selection in the genome. This has important implications to empirical estimates of *r*_*mf*,*W*_, especially in the context of recent theoretical work (Flintham *et al*., 2023).

Traditional population genetic models of SA selection predict that stable SA polymorphisms are possible only if selection coefficients in males and females are large (Kidwell *et al*., 1977) - unless SA loci exhibit sex-specific dominance reversals (Fry, 2010). It has been argued that if sex-specific fitness landscapes are concave (or decelerating) in the vicinity of fitness optima, sex-specific dominance patterns at SA loci can emerge naturally such that the allele that benefits females is dominant in females and the allele that benefits males is dominant in males. Yet, detecting SA loci has proved to be notoriously difficult (Ruzicka *et al*., 2020). In a recent study, Flintham *et al*. (2023) pointed out that most traditional models investigating the maintenance of SA genetic polymorphisms investigated only one or two biallelic loci. In reality, traits under SA selection are likely to be polygenic in nature (Pennell *et al*., 2024). Flintham *et al*. (2023) considered two different genetic architectures for a trait perfectly genetically correlated between the sexes but experiencing SA selection: (1) a continuum of alleles model where the trait evolves through recurrent occurrence and (possible) fixation of new additive mutations at a single diploid locus, and (2) a polygenic model where the trait evolves as a result of evolution at an arbitrary number of diallelic loci. Their mathematical treatment as well as individual based simulations suggest that SA selection can facilitate the maintenance of genetic polymorphisms only when selection is “diversifying”, i.e., when fitness functions are accelerating and the strength of directional selection is sufficiently strong. Flintham *et al*. (2023) argued that such a delicate balance is very unlikely to be realised in nature. When fitness functions are decelerating (or concave), polymorphisms are maintained at - at the most - one locus, irrespective of the number of loci coding for the trait. Furthermore, their multilocus simulations highlight that even such SA balanced polymorphisms are rather transient on evolutionary timescales, and therefore their effects are unlikely to be detected at genomic scales. In the light of these theoretical predictions, it is unclear whether negative empirical estimates of *r*_*mf*,*W*_ (Chippindale *et al*., 2001; Delcourt *et al*., 2009; Innocenti & Morrow, 2010) reflect true SA polymorphisms. Our findings suggest that these negative estimates of *r*_*mf*,*W*_ could alternatively be a result of negative LD between female-limited and male-limited loci.

Our simulations also exhibit considerable variation in *r*_*mf*,*W*_ between replicates of our simulations. For example, for the case when the map length of the simulated genome was 0.1 M and the mutation rate was 7 × 10^*−*7^, *r*_*mf*,*W*_ for independent replicate simulations vary between approximately -0.300 to 0.125. Since direct selective effects do not contribute in any way to *r*_*mf*,*W*_ in the purely sex-limited selection scenario we simulated, this variation in *r*_*mf*,*W*_ is driven by indirect selective effects alone - specifically by variation in average effects at various male-limited and female-limited loci and the LD between them. If the patterns of LD between male-limited and female-limited loci evolve rapidly due to genetic drift, signals of sexual antagonism would vary greatly even among closely related populations of the same species. Collet *et al*. (2016) measured *r*_*mf*,*W*_ in two replicate of the “*LH*_*M*_ “ laboratory population of *D. melanogaster* maintained in two different laboratories. They found that *r*_*mf*,*W*_ was negative in one of the replicates, but not significantly different from 0 in the other. Interestingly, a more recent investigation on yet another replicate of the “*LH*_*M*_ “ population in a third laboratory has reported a significantly positive estimate of *r*_*mf*,*W*_ (Geeta Arun *et al*., 2022). Collet *et al*. (2016) argued that genetic drift was unlikely to be the cause of the between-population divergence in *r*_*mf*,*W*_, since they found non-random patterns associated with allele frequency change that were not consistent with drift. However, genetic drift can also result in the evolution of the linkage structure of the population affecting the contribution made by indirect selection to *COV*_*mf*,*W*_ (second term of Equation 1). Therefore, in principle, differences in estimates of *r*_*mf*,*W*_ from closely related populations of the same species could just be a by-product of differences in the LD structure of those populations.

### 6.2 Purely sex-limited selection can result in selective interference between the sexes

In its classical formalism, Intralocus Sexual Conflict (IaSC) occurs when a trait with a common genetic basis between the sexes experiences SA selection (Bonduriansky & Chenoweth, 2009; Cox & Calsbeek, 2009; Van Doorn, 2009). This evolutionary “tug-of-war” between the sexes is thought to lead to the evolution of sex-specific genetic architectures allowing the sexes to evolve to their respective fitness optima resulting in the evolution of sexual dimorphism (Lande, 1980; Rice, 1984; Reeve & Fairbairn, 2001; Pennell *et al*., 2024). Often, sex-biased gene-expression is considered as evidence of resolved IaSC (Innocenti & Morrow, 2010; Jaquiéry *et al*., 2013). Our results suggest that even when the genetic architecture for fitness is perfectly sex-limited, HRI can result in selective interference between the sexes. While a negative *r*_*mf*,*W*_ suggests selective interference between the sexes, it does not provide any information on the magnitude of this effect in the context of rates of adaptation. To understand the magnitude of such HRI-driven selective interference between the sexes we make use of Fisher’s Fundamental Theorem of Natural Selection (FTNS) (Queller, 2017; Walsh & Lynch, 2018). FTNS shows that the rate of change of mean absolute fitness of a population due to one generation of natural selection - in the absence of mutation and migration - is exactly equal to the additive genetic variance for relative fitness in that population. FTNS can be readily generalised to a case where selection operates in different ways in males and females (Martinossi-Allibert *et al*., 2018), and can be described as follows:

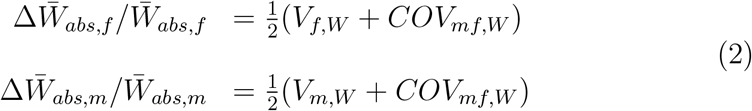

where 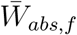 and 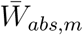 are the mean absolute fitnesses in females and males respectively. Sex-specific FTNS suggests that the rate of adaptation in one sex is equal to the arithmetic mean of the additive genetic variance for relative fitness in that sex and the intersexual additive genetic covariance for relative fitness. Notice that if *COV*_*mf*,*W*_ is negative selection in one sex impedes adaptation in the other sex. The strength of this selective interference between the sexes depends on the relative magnitudes of *V*_*f*,*W*_, *V*_*m*,*W*_, and *COV*_*mf*,*W*_ . Therefore, we measured the percentage change in the rate of adaptation in females due to this effect as 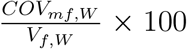. We found that if the recombination rate is low, or if the mutation rate is high, HRI can result in nearly 10-15% retardation in the rate of adaptation in females. It is important to note that this is the median of our replicate simulations. For a substantial fraction of the simulations the retardation of the rate of adaptation in females was even stronger.

### 6.3 Conclusion

In a recent review Pennell *et al*. (2024) argued against the use of the phrase “Intralocus Sexual Conflict”. They noted that sex-specific selection involves an interplay of many different fitness-related traits - and therefore, sex-specific selection is best understood as a multi-locus problem. Flintham *et al*. (2023) analysed poly-allelic and polygenic models of sex-specific selection and derived results that run counter to the conventional wisdom derived from single-locus population genetic models. Complementary to these studies, our study underlines the fact that the moment one considers sex-specific fitness as a multilocus trait one has to grapple with important multilocus phenomena such as Hill-Robertson Interference between fitness-determining loci. Our results highlight the importance of taking into consideration indirect selective effects mediated by LD between fitness-determining loci. We show that such indirect selection coupled with HRI can drive *r*_*mf*,*W*_ to be negative even in the absence of sexually antagonistic selection. Given that SA selection is neither necessary (our results) nor sufficient (results by Connallon & Matthews (2019)) for *r*_*mf*,*W*_ to be negative, it is clear that *r*_*mf*,*W*_ has limited utility as a measure of the strength of IaSC. We argue that the relative magnitude the selective interference between the sexes given by 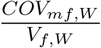 or 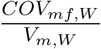 is a more useful measure of the strength of selective interference between the sexes. Finally, our results show that contrary to canonical descriptions of IaSC, even fully sex-limited genetic architectures can result in selective interference between the sexes.

## 7 Acknowledgments

Computer simulations and downstream analyses described here were performed using the AC3 computing cluster facility based at Ashworth Laboratories, the University of Edinburgh. This work was supported by a Natural Environmental Research Council (NERC, UK) grant (project number “10568701 10568704”) to Jarrod Hadfield and Darren Obbard. MGA thanks Soumya Panyam and Broti Biswas for their feedback on an earlier version of this manuscript.

## 8 Supplementary Figures

**Figure S1:**
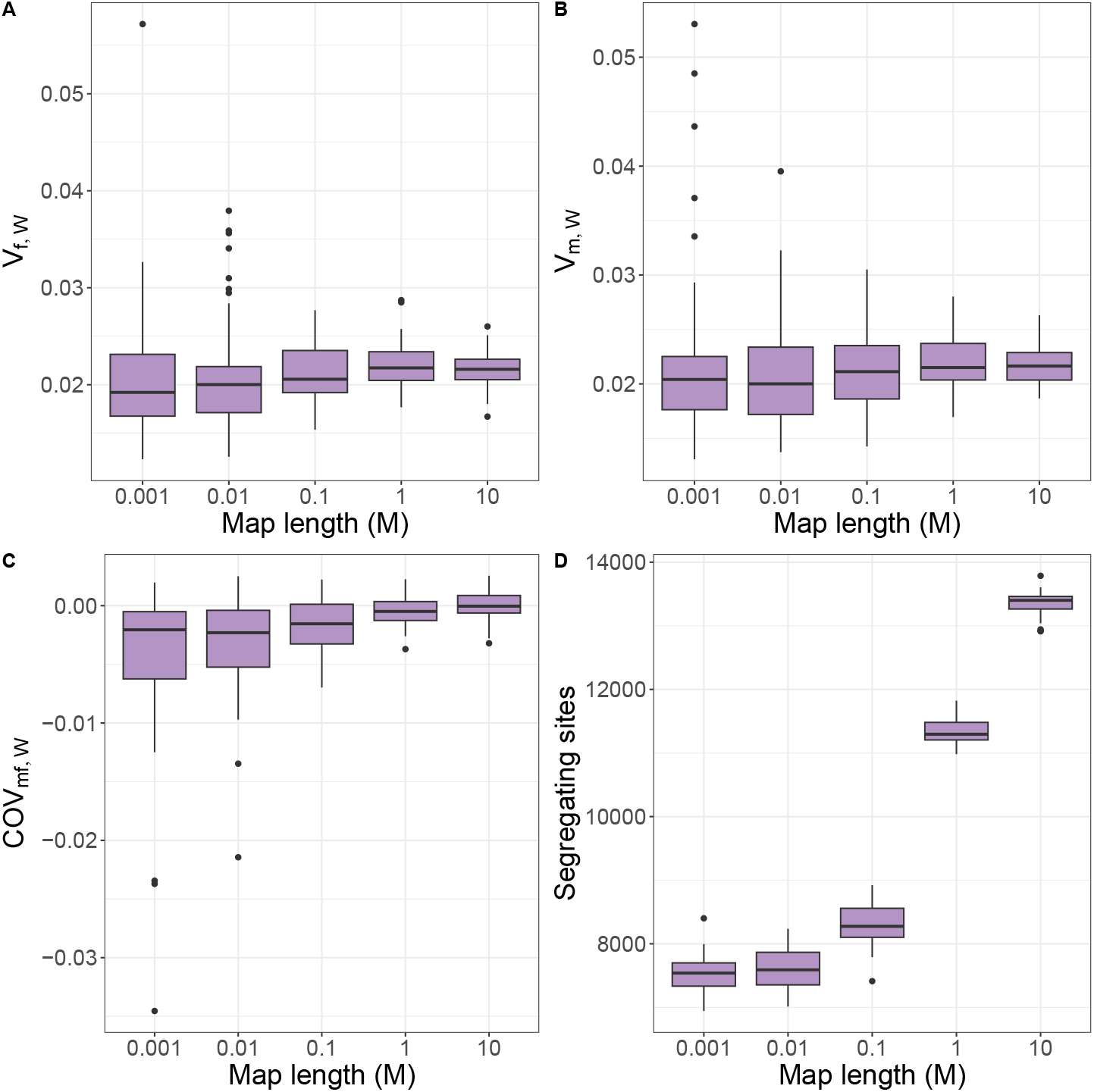
Boxplots representing (A) additive genetic variance for female relative fitness (*V*_*f*,*W*_), (B) additive genetic variance for male relative fitness (*V*_*m*,*W*_), (c) intersexual additive genetic covariance for relative fitness (*COV*_*mf*,*W*_), and (D) the number of segregating sites in the population as a function of the map length of the simulated region. Each boxplot is based on 50 replicate simulations. In these simulations the mutation rate was fixed at 7 × 10^*−*7^ per sit per genome.

**Figure S2:**
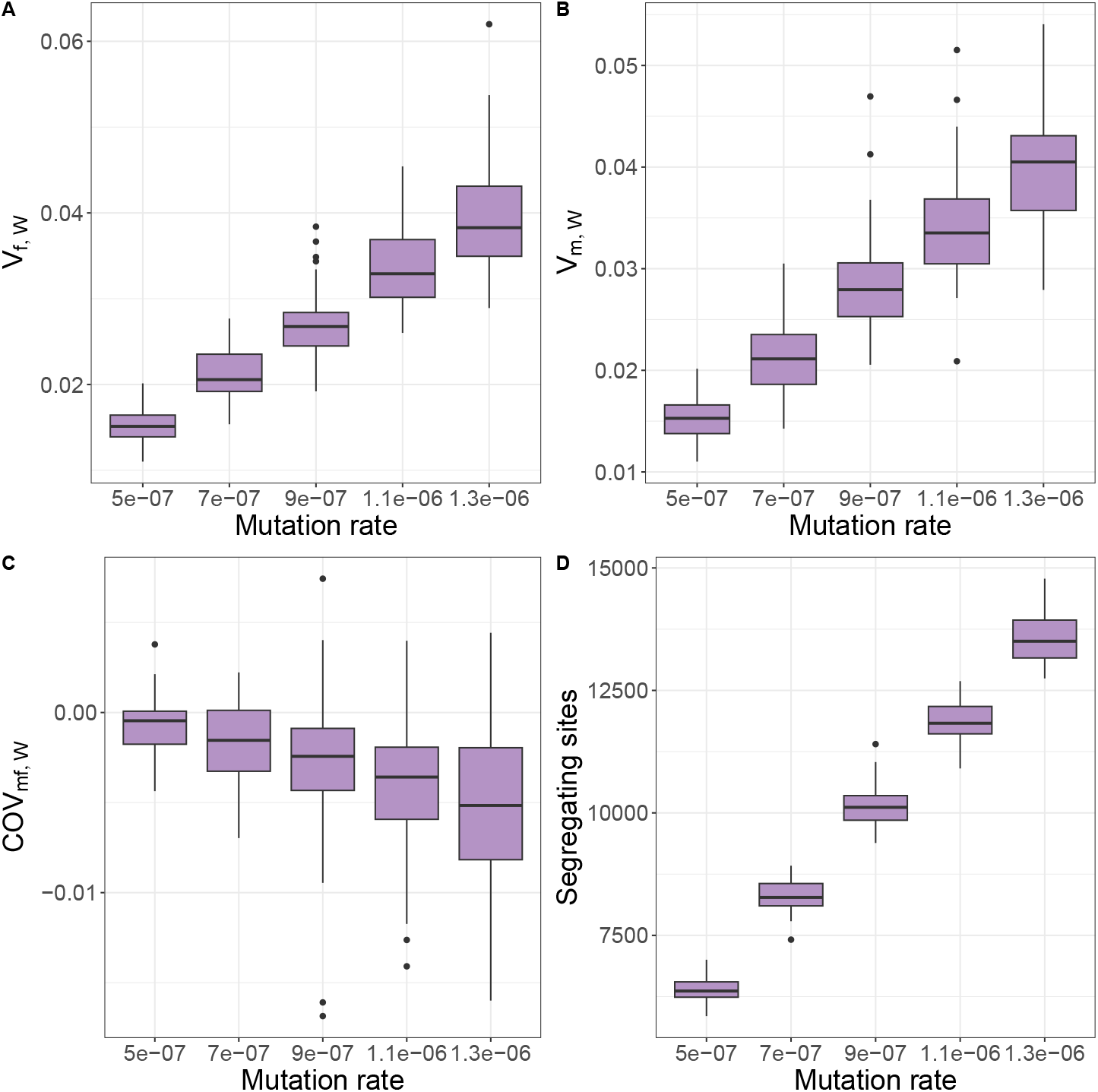
Boxplots representing (A) additive genetic variance for female relative fitness (*V*_*f*,*W*_), (B) additive genetic variance for male relative fitness (*V*_*m*,*W*_), (c) intersexual additive genetic covariance for relative fitness (*COV*_*mf*,*W*_), and (D) the number of segregating sites in the population as a function of the mutation rate. Each boxplot is based on 50 replicate simulations. In these simulations the map length of the simulated region was fixed at 0.1 M.

## References

Berger, D., Grieshop, K., Lind, M.I., Goenaga, J., Maklakov, A.A. & Arnqvist, G. (2014). Intralocus sexual conflict and environmental stress. Evolution, 68, 2184–2196.

Berger, D., Martinossi-Allibert, I., Grieshop, K., Lind, M.I., Maklakov, A.A. & Arnqvist, G. (2016). Intralocus sexual conflict and the tragedy of the commons in seed beetles. The American Naturalist, 188, E98–E112.

Bonduriansky, R. & Chenoweth, S.F. (2009). Intralocus sexual conflict. Trends in ecology & evolution, 24, 280–288.

Buffalo, V. & Coop, G. (2019). The linked selection signature of rapid adaptation in temporal genomic data. Genetics, 213, 1007–1045.

Buffalo, V. & Coop, G. (2020). Estimating the genome-wide contribution of selection to temporal allele frequency change. Proceedings of the National Academy of Sciences, 117, 20672–20680.

Chippindale, A.K., Gibson, J.R. & Rice, W.R. (2001). Negative genetic correlation for adult fitness between sexes reveals ontogenetic conflict in drosophila. Proceedings of the National Academy of Sciences, 98, 1671–1675.

Collet, J.M., Fuentes, S., Hesketh, J., Hill, M.S., Innocenti, P., Morrow, E.H., Fowler, K. & Reuter, M. (2016). Rapid evolution of the intersexual genetic correlation for fitness in drosophila melanogaster. Evolution, 70, 781–795.

Collet, J.M. & Sztepanacz, J.L. (2022). Mating traits are phenotypically but not genetically correlated to fitness. Behavioral Ecology, 33, 833–843.

Connallon, T. & Clark, A.G. (2010). Sex linkage, sex-specific selection, and the role of recombination in the evolution of sexually dimorphic gene expression. Evolution, 64, 3417–3442.

Connallon, T. & Matthews, G. (2019). Cross-sex genetic correlations for fitness and fitness components: connecting theoretical predictions to empirical patterns. Evolution letters, 3, 254–262.

Cox, R.M. & Calsbeek, R. (2009). Sexually antagonistic selection, sexual dimorphism, and the resolution of intralocus sexual conflict. The American Naturalist, 173, 176–187.

Day, T. & Bonduriansky, R. (2004). Intralocus sexual conflict can drive the evolution of genomic imprinting. Genetics, 167, 1537–1546.

Delcourt, M., Blows, M.W. & Rundle, H.D. (2009). Sexually antagonistic genetic variance for fitness in an ancestral and a novel environment. Proceedings of the Royal Society B: Biological Sciences, 276, 2009–2014.

Delph, L.F., Andicoechea, J., Steven, J.C., Herlihy, C.R., Scarpino, S.V. & Bell, D.L. (2011). Environment-dependent intralocus sexual conflict in a dioecious plant. New Phytologist, 192, 542–552.

Fedorka, K.M. & Mousseau, T.A. (2004). Female mating bias results in conflicting sex-specific offspring fitness. Nature, 429, 65–67.

Felsenstein, J. (1974). The evolutionary advantage of recombination. Genetics, 78, 737–756.

Flintham, E., Savolainen, V., Otto, S., Reuter, M. & Mullon, C. (2023). The maintenance of genetic polymorphism in sexually antagonistic traits. bioRxiv, pp. 2023–10.

Foerster, K., Coulson, T., Sheldon, B.C., Pemberton, J.M., Clutton-Brock, T.H. & Kruuk, L.E. (2007). Sexually antagonistic genetic variation for fitness in red deer. Nature, 447, 1107–1110.

Fry, J.D. (2010). The genomic location of sexually antagonistic variation: some cautionary comments. Evolution, 64, 1510–1516.

Geeta Arun, M., Chechi, T.S., Meena, R., Bhosle, S.D. Srishti & Prasad, N.G. (2022). Investigating the interaction between inter-locus and intralocus sexual conflict using hemiclonal analysis in drosophila melanogaster. BMC Ecology and Evolution, 22, 38.

Haldane, J. (1962). Conditions for stable polymorphism at an autosomal locus. Nature, 193, 1108–1108.

Haller, B.C. & Messer, P.W. (2023). Slim 4: multispecies eco-evolutionary modeling. The American Naturalist, 201, E127–E139.

Hill, W.G. & Robertson, A. (1966). The effect of linkage on limits to artificial selection. Genetics Research, 8, 269–294.

Innocenti, P. & Morrow, E.H. (2010). The sexually antagonistic genes of drosophila melanogaster. PLoS biology, 8, e1000335.

Jaquiéry, J., Rispe, C., Roze, D., Legeai, F., Le Trionnaire, G., Stoeckel, S., Mieuzet, L., Da Silva, C., Poulain, J., Prunier-Leterme, N. et al. (2013). Masculinization of the x chromosome in the pea aphid. PLoS Genetics, 9, e1003690.

Kappeler, P.M., Benhaiem, S., Fichtel, C., Fromhage, L., Höner, O.P., Jennions, M.D., Kaiser, S., Krüger, O., Schneider, J.M., Tuni, C. et al. (2023). Sex roles and sex ratios in animals. Biological Reviews, 98, 462–480.

Kidwell, J., Clegg, M., Stewart, F. & Prout, T. (1977). Regions of stable equilibria for models of differential selection in the two sexes under random mating. Genetics, 85, 171–183.

Lande, R. (1980). Sexual dimorphism, sexual selection, and adaptation in polygenic characters. Evolution, pp. 292–305.

Long, T.A. & Rice, W.R. (2007). Adult locomotory activity mediates intralocus sexual conflict in a laboratory-adapted population of drosophila melanogaster. Proceedings of the Royal Society B: Biological Sciences, 274, 3105–3112.

Martinossi-Allibert, I., Savković, U., orević, M., Arnqvist, G., Stojković, B. & Berger, D. (2018). The consequences of sexual selection in well-adapted and maladapted populations of bean beetles. Evolution, 72, 518–530.

Owen, A. (1953). A genetical system admitting of two distinct stable equilibria under natural selection. Heredity, 7, 97–102.

Parsons, P. (1961). The initial progress of new genes with viability differences between sexes and with sex linkage. Heredity, 16, 103–107.

Pennell, T.M., Mank, J.E., Alonzo, S.H. & Hosken, D.J. (2024). On the resolution of sexual conflict over shared traits. Proceedings B, 291, 20240438.

Price, D.K. & Burley, N.T. (1993). Constraints on the evolution of attractive traits: genetic (co) variance of zebra finch bill colour. Heredity, 71, 405–412.

Queller, D.C. (2017). Fundamental theorems of evolution. The American Naturalist, 189, 345–353.

Reeve, J. & Fairbairn, D. (2001). Predicting the evolution of sexual size dimorphism. Journal of Evolutionary Biology, 14, 244–254.

Rice, W.R. (1984). Sex chromosomes and the evolution of sexual dimorphism. Evolution, pp. 735–742.

Roze, D. & Barton, N.H. (2006). The hill–robertson effect and the evolution of recombination. Genetics, 173, 1793–1811.

Ruzicka, F., Dutoit, L., Czuppon, P., Jordan, C.Y., Li, X.Y., Olito, C., Runemark, A., Svensson, E.I., Yazdi, H.P. & Connallon, T. (2020). The search for sexually antagonistic genes: Practical insights from studies of local adaptation and statistical genomics. Evolution letters, 4, 398–415.

Ruzicka, F., Hill, M.S., Pennell, T.M., Flis, I., Ingleby, F.C., Mott, R., Fowler, K., Morrow, E.H. & Reuter, M. (2019). Genome-wide sexually antagonistic variants reveal long-standing constraints on sexual dimorphism in fruit flies. PLoS Biology, 17, e3000244.

Singh, A., Hasan, A. & Agrawal, A.F. (2023). An investigation of the sex-specific genetic architecture of fitness in drosophila melanogaster. Evolution, 77, 2015–2028.

Spencer, H. & Clark, A. (2014). Non-conflict theories for the evolution of genomic imprinting. Heredity, 113, 112–118.

Spencer, H.G. & Priest, N.K. (2016). The evolution of sex-specific dominance in response to sexually antagonistic selection. The American Naturalist, 187, 658–666.

Van Doorn, G.S. (2009). Intralocus sexual conflict. Annals of the New York Academy of Sciences, 1168, 52–71.

Vincent, C.M. & Sharp, N.P. (2014). Sexual antagonism for resistance and tolerance to infection in drosophila melanogaster. Proceedings of the Royal Society B: Biological Sciences, 281, 20140987.

Walsh, B. & Lynch, M. (2018). Evolution and selection of quantitative traits. Oxford University Press.

Wolak, M.E., Arcese, P., Keller, L.F., Nietlisbach, P. & Reid, J.M. (2018). Sex-specific additive genetic variances and correlations for fitness in a song sparrow (melospiza melodia) population subject to natural immigration and inbreeding. Evolution, 72, 2057–2075.

Wong, H.W. & Holman, L. (2023). Pleiotropic fitness effects across sexes and ages in the drosophila genome and transcriptome. Evolution, 77, 2642–2655.

